# A controlled thermoalgesic stimulation device to identify novel pain perception biomarkers

**DOI:** 10.1101/2020.07.01.177568

**Authors:** Maider Núñez Ibero, Borja Camino-Pontes, Ibai Diez, Asier Erramuzpe, Endika Martínez Gutiérrez, Sebastiano Stramaglia, Javier Ortiz Álvarez-Cienfuegos, Jesus M. Cortes

## Abstract

**Objective:** To develop a new device that will help identify physiological markers of pain perception by reading the brain’s electrical activity and the bodies hemodynamic interactions while applying thermoalgesic stimulation. Methods: We designed a compact prototype that generates well-controlled thermal stimuli using a computer driven Peltier cell while simultaneously capturing electroencephalography (EEG) and photoplethysmography (PPG) signals as the stimuli are varied. The study was performed on 35 healthy subjects (mean age 30.46 years, SD 4.93 years; 20 males, 15 females) and to account for the inter-subject variability in the tolerance to thermal pain, we first determined the heat pain threshold (HPT) for each subject, defined as the maximum temperature that the subject can withstand when the Peltier cell gradually increases the temperature. Subsequently, we defined the pain parameters associated with a stimulation temperature equivalent to 90% of the HPT, comparing this to the no-pain state (control) in the absence of thermoalgesic stimulation. Results: Both the one-dimensional and the two-dimensional spectral entropy (SE) obtained from both the EEG and PPG signals could differentiate the condition of pain. In particular, the PPG SE was significantly reduced in association with pain, while the SE for EEG increased slightly. Moreover, significant discrimination occurred within a specific range of frequencies, 26-30 Hz for EEG and about 5-10 Hz for PPG. Conclusion: Hemodynamics, brain dynamics and their interactions can discriminate thermal pain perception. Significance: The possibility of monitoring on-line variations in thermal pain perception using a similar device and algorithms may be of interest to study different pathologies that affect the peripheral nervous system, such as small fiber neuropathies, fibromyalgia or painful diabetic neuropathy.

## I. Introduction

**T**HE synergy between electronic technology and state-of-the-art instrumentation, together with the incorporation of statistical analysis and data science, provides tremendous possibilities in neuroscience research [1]. Here we have designed a new, compact hardware-device to measure systemic responses of the peripheral nervous system (PNS), such as the perception of pain, by progressively increasing a Peltier cell’s temperature in contact with a subjects skin or hand. The device allows simultaneously recording of brain and heart dynamics by measuring electroencephalography (EEG) and photoplethysmography (PPG)^1^, respectively. Therefore, our device can be used to quantitatively measure systemic physiological responses to thermal stimulation, allowing the underlying structural and functional changes to be elucidated, and offering an insight into the physiological interactions provoked, the so-called physiolome [2]–[4].

But what exactly is pain perception and how can it be measured? A definition of pain was formulated more than 50 years ago [5]: *Pain is an unpleasant experience that we primarily associate with tissue damage or describe in terms of tissue damage or both.* Since then, multidisciplinary approaches and the emergence of models for chronic pain-related disease have produced substantial advances in our understanding of pain, its assessment and treatment. As such, a more refined IASP’s definition has been proposed, whereby: *Pain is an unpleasant sensory and emotional experience associated with actual or potential tissue damage, or described in terms of such damage*. Accordingly, it is now well-accepted that pain encompasses a systemic response that can be detected or perceived over quite different scales and systems.

Here we have asked whether pain perception might be encoded through different physiological signals and we sought to assess their possible interactions. The physiological response to pain has been addressed previously using approaches like EEG [6], [7] and PPG [8], [9], yet these signals are typically analyzed separately. Moreover, the paradigm to produce painful stimulation relies on human intervention [10] or environmental factors [11], which may compromise the reliability of these results. By contrast, the device we have designed produces well-controlled painful stimulation. Although not yet approved by the Food and Drug Administration (FDA), General Electric Healthcare introduced the Surgical Pleth Index (SPI) to measure the increase in sympathetic activity from the PPG signal in response to painful (nociceptive) stimuli [12]. However, the SPI only works in conjunction with general anesthesia and thus, much of the cortical processing that occurs when a painful stimulus is received will be ignored. Moreover, the precise relationship between the entropy of physiological responses and pain perception remains unclear. Nevertheless, the relationship between variations in entropy following exposure to a nociceptive stimulus has been assessed previously [13], showing an increase in the spatial entropic patterns in response to painful stimuli.

Here we have studied the dynamic physiological interactions that are produced in response to a painful stimulus in an attempt to define the perception of pain. In contrast to other studies, we designed and used a device that objectively controls the painful thermal stimulus generated, whilst synchronously recording electrical neuronal activity and some hemodynamic parameters. The main working hypothesis was that by simultaneously monitoring these variables in response to a painful stimulus and comparing them to the basal response, novel aspects of sympathetic excitation and regulation will be revealed that are related to the perception of pain. From the data obtained, we intend to validate the usefulness of our system to evaluate groups of patients with different pathologies associated with abnormalities in the PNS.

## II. Methods

### A. Hardware

To design and manufacture an electronic device to generate the stimuli, we used OrCAD (version 16.6 Lite) to automate the production of the printed circuit, the board design and photolithography, and for the chemical etching to finally manufacture the boards. Once the circuit boards were designed and manufactured, the electronic components were inserted and soldered onto them at the Electronic Technology laboratory in the Bilbao School of Engineering. We used MATLAB (version R2017a, MathWorks Inc., Natick, MA, USA) to create the user interface, connecting it to the hardware device that generates the stimuli, and to process the physiological signals, run the spectral entropy (SE) algorithms, presents the results and prepare the final images. The BioPac system (BioPac Systems, Inc, Student Lab MP36) was configured to read the physiological variables and the data obtained was processed using the AcqKnowledge software (version BSL PRO 3.7), which records, analyses and filters the data in real-time, presenting it as a continuous record, an X-Y chart or a histogram.

### B. Participants and Ethical considerations

The study was carried out on 35 healthy volunteers (20 men; 15 women) recruited at the University of the Basque Country and with a mean age of 30.46 years (SD 4.93 years). All the participants provided their signed informed consent and the study was approved by the Ethical Committee of the University of the Basque Country (project 2017/092). The data were acquired according to the guidelines laid down by the University’s Ethical Committee and the Ethical Principles for Medical Research Involving Human Subjects set out in the Helsinki Declaration. The inclusion criteria were to be aged between 20 and 40 years-old and to have provided signed informed consent. The exclusion criteria were any diagnosed illness, medication use or drug consumption in the month prior to testing, the refusal of the volunteer to participate in the study or the consumption of energetic drinks immediately prior to testing.

### C. Capture and cleaning of physiological signals

Before starting the experiments, the subjects were seated near a table with a computer on it, listening to relaxing music so that the experiments commenced in a calm and emotional state. In order to minimize eye-blinking artifacts, all subjects were asked to remain silent with their eyes closed, and to stay still without crossing their legs throughout the experiment. The physiological variables captured were the EEG signals from both cerebral hemispheres and the PPG signals from the contralateral hand to that being heated by the Peltier cell (see figure 1A).

**Fig. 1.**
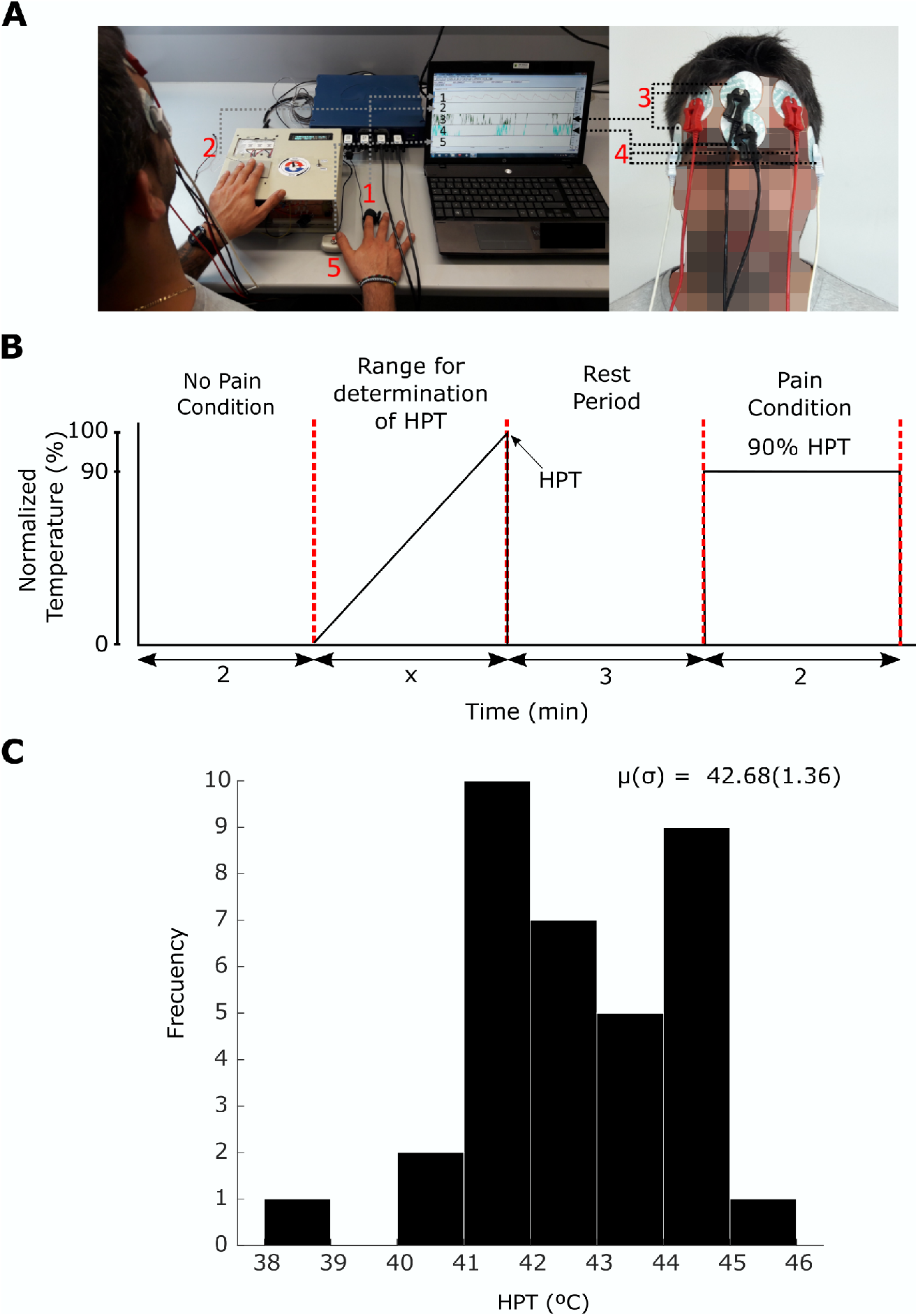
Device for pain perception studies. **A**: System for the capture and recording of physiological signals. The PPG sensor (1) is attached to the distal phalanx of the index finger of the subject’s hand, while the opposite hand is placed on the Peltier cell (2). The electrodes for the EEG signals (3,4) are placed on both sides of the forehead. Finally, the push-button is used to determine the thermoalgesic threshold. **B**: Pain stimulation is divided into four distinct regions, the first lasts 2 min in the absence of any painful stimulation to define the no pain condition. The second focuses on calculating the heat pain threshold (HPT) of *x* min, which was different for each participant. This phase was achieved by increasing the temperature in increments of 0.5°C up to the HPT, the maximum temperature that the subject withstood. After a rest period of 3 min, a final regime to define the pain condition was achieved at the temperature equivalent to 90% of the HPT, lasting for 2 min. **C**: HPT histogram of all the participants, indicating the mean *μ* and standard deviation *σ*.

The EEG signals were obtained through six electrodes situated in accordance with the Electrode Position Modified Combinatorial Nomenclature [14], two on one cerebral hemisphere, another two on the other hemisphere and two more as reference points. The electrodes used were silver/silver chloride gel types. The EEG electrodes were located on the forehead using a bipolar arrangement, measuring the differentiated voltages between the FP1 (Frontal Pole) and AF7 (Anterior Frontal) electrodes in one of the channels, and between the symmetrical FP2 and AF8 points in the second channel. For the reference electrodes, the first channel uses an electrode in the lower front central position, while the second channel uses the upper front central position. A differential distribution was employed with separate but analogous reference points, such that the readings from the left and right hemispheres are largely comparable. The EEG electrode location at the frontal positions guarantees a superior amplitude and integrity of the signals acquired, resulting in a lower impedance of the electrode-skin interface. Moreover, the prefrontal locations of the EEG might modulate the pain-related autonomic response [15].

The PPG signal was acquired by attaching a transducer to the tip of the index finger, consisting of a matched infrared emitter and a photodiode detector that transmits the changes in infrared reflectance resulting from the variation in blood flow. When the PPG transducer is placed on the skin, close to the capillaries, the reflectance of the infrared light from the emitter to the detector will change in accordance with the capillary blood volume, enabling the blood volume pulse waveform to be recorded. It is important to note that the PPG signal is an efficient and interesting alternative to measure heartbeat intervals, since it is simpler than the electrocardiogram while achieving a precise measurement for heart rate variability [16].

All the signals were filtered with a 38.5 Hz Low Pass Notch Filter incorporated into the series amplifier modules of BioPac, thereby eliminating the 50 Hz mains interference. Two differentiated signals (FP1 minus AF7 and FP2 minus AF8) were obtained and filtered using a recent data-driven algorithm that removes ocular and muscle artifacts from the single-channel data, referred to as the surrogate-based artifact removal (SuBAR) method [17]. Although the full details are given elsewhere [17], the algorithm follows the pipeline: 1, Z-score of the data; 2, Maximal Overlap Wavelet Transform (MODWT) of the data, using symlets of order 5 and with 5 levels of decomposition; 3, Removal of artifacts, defined as the values of the wavelet coefficients that are outliers relative to the distribution of the values obtained from the data surrogates (to identify outliers, we considered a 5% significance level and the outlier coefficients were eliminated by substituting their values with the average coefficients obtained from the surrogates); 4. Reconstruction of the time-domain signal using the inverse MODWT and the cleaned coefficients. All the calculations were implemented in MATLAB (version R2019a, MathWorks Inc., Natick, MA, USA).

### D. Stimulation protocol

The system for data capture is shown in Figure 1A. There are two main elements to this set-up, the BioPac and the equipment to generate the stimulus connected to a portable unit. The AcqKnowledge program displays the signals received from the PPG sensor (1), the sensor temperature (2) and the EEG (3,4), as well as the reference points inserted by a push-button to mark the point the heat pain threshold is reached (5), the maximum temperature that the subject can withstand in terms of exposure to heat pain. The no-pain condition (control) was obtained in complete absence of a painful stimulation (figure 1B), measuring the electrophysiological signals while keeping the participant’s hand off the Peltier cell. This condition persisted for two minutes.

Signals were recorded over another two or three minutes to estimate the heat pain threshold (HPT) [18]. This was calculated while the participant placed one hand on the Peltier cell, recording the variables using the BioPac and progressively increasing the cell’s temperature in a controlled fashion, varying it in increments of 0.5°C up to the maximum temperature that the participant withstood, the HPT. This value was subject-dependent. The strategy of progressively increasing the temperature was critical in these experiments and it was performed in this way to achieve dual activation of the C receptors responsible for heat sensing and the A*δ* fiber receptors that process noxious stimuli [19], [20].

A 2 minute recording was obtained during the pain stimulus, defined at a temperature equal to 90% of the HPT and chosen in this way as the maximum as possible without exceeding the Ethical Committee’s recommendations. All the temperature values were transformed to values relative to the HPT and this normalization permits a comparative analysis across subjects.

### E. Spectral entropy to detect pain perception

The SE is a generalization of the Shannon entropy, where the state probability *p*(*x*) is replaced by a normalized power spectral density *p*(*f*), which represents the probability density function of the power as a function of frequency. Here, the power SE was calculated through the absolute square of the Fast Fourier Transformation, calculated with the function *fft* in MATLAB. After normalization, we obtained *p*(*f*) and from there, the one-dimensional spectral entropy was calculated as:

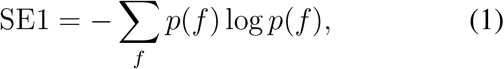

and applied individually to the three signals: EEG1 (left hemisphere), EEG2 (right hemisphere), and PPG.

For the two-dimensional SE we made use of the two-dimensional Fast Fourier *fft2* transformation in MATLAB, and after normalization, we defined:

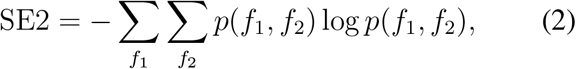

applied to any pairs of variables in the triplet EEG1, EEG2 and PPG.

Here, both SE1 and SE2 were calculated in 80%-of-overlapping windows of 5 Hz, and the code can be downloaded at https://github.com/compneurobilbao/spectral-entropy-maider.

### F. Statistical Analyses

For both SE1 and SE2, we calculated the FFT using the Blackman window function (implemented as *blackman* in Matlab) over time windows of 500 time points, and with a sampling frequency of 500 Hz, corresponding to 1 sec. After sliding the time window, we obtained a time series for SE1 and SE2, the latter representing a temporal sequence of the SE2 matrices. For each frequency value, the final SE1 and SE2 values were obtained by averaging all the entropy values in the temporal dimension. These temporal mean values of SE1 and SE2 were those used for the statistical comparison between the pain and no-pain conditions.

The discriminability between conditions was calculated as 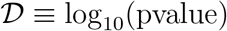 and four stages were followed to obtain the p-values. We first calculated the SE1 and SE2 time series for each condition and subject, and we then averaged the two metrics over the entire time dimension. We then employed a Wilcoxon signed-rank test between the conditions, with the different subject measures considered as observations. Finally, to correct for multiple comparisons we applied a false-discovery-rate (FDR) and Bonferroni corrections, the latter using a significance threshold equal to 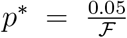, where 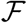 corresponds to the number of different frequencies used to compare the SE values (40 EEG and 10 for PPG). All the statistical analyses were performed in MATLAB (version R2019a, MathWorks Inc., Natick, MA, USA).

## III. Results

A cohort of N = 35 subjects participated in the study and we obtained a HPT value for each subject when stimulated with our device (figure 1C, Table I), with a mean HPT of 42.68°C (SD 1.36). There were apparently no significant gender differences in the HPT, as witnessed when the HPT of a subgroup of males (N=16, mean age 30.04 years: t-test=-0.46, p-value=0.64) was compared with an age matched group of females (N=16, mean age 29.23 years: t-test=-0.45, p-value=0.63). This result was consistent with previous studies showing no differences over an age range similar to ours [21]. Importantly, normalization of the temperature values to the HPT allowed the two conditions of pain and no-pain to be defined independently of the participant, making the different metrics across participants and conditions comparable.

**TABLE I.** Sex, age, hand stimulated and Heat Pain Threshold (°C) for all participants (N=35, 20 men;15 women)

|  | Sex | Age | Hand Heated | Heat Pain Threshold *HPT* |
| --- | --- | --- | --- | --- |
| Subject 1 | M | 32 | Right | 43.23 |
| " 2 | F | 28 | Right | 45.02 |
| " 3 | M | 24 | Right | 41.23 |
| " 4 | F | 30 | Right | 44.12 |
| " 5 | M | 30 | Right | 43.00 |
| " 6 | M | 33 | Right | 42.75 |
| " 7 | M | 36 | Right | 38.80 |
| " 8 | M | 33 | Right | 41.89 |
| " 9 | F | 36 | Right | 42.13 |
| " 10 | F | 31 | Right | 41.60 |
| " 11 | F | 31 | Right | 44.42 |
| " 12 | M | 23 | Right | 44.03 |
| " 13 | F | 30 | Right | 43.47 |
| " 14 | M | 31 | Right | 40.98 |
| " 15 | F | 40 | Left | 42.82 |
| " 16 | F | 20 | Right | 42.60 |
| " 17 | M | 20 | Right | 44.67 |
| " 18 | M | 38 | Right | 41.22 |
| " 19 | M | 26 | Right | 42.32 |
| " 20 | F | 25 | Right | 42.49 |
| " 21 | F | 31 | Right | 43.07 |
| " 22 | M | 30 | Right | 42.89 |
| " 23 | M | 36 | Right | 44.10 |
| " 24 | M | 36 | Right | 44.23 |
| " 25 | M | 31 | Left | 44.10 |
| " 26 | M | 31 | Left | 41.52 |
| " 27 | F | 24 | Left | 43.10 |
| " 28 | F | 30 | Left | 41.35 |
| " 29 | M | 33 | Left | 41.56 |
| " 30 | F | 27 | Left | 41.84 |
| " 31 | M | 33 | Left | 44.10 |
| " 32 | M | 28 | Left | 44.3 |
| " 33 | F | 22 | Left | 40.95 |
| " 34 | M | 35 | Left | 41.83 |
| " 35 | F | 26 | Left | 41.91 |

The SE values obtained from the different physiological signals between the pain and no-pain conditions were compared following the procedure explained in figure 2. The statistical significance of all the possible comparisons was assessed with the discriminability (D) obtained from the p-values after a Wilcoxon signed-rank test was performed on the data from the different conditions (Methods, the discriminability is illustrated in figure 3A for the 1D case). Initially, the SE1 time series was calculated for each condition and participant, and the temporal averages were then calculated. These values were compared between conditions and across the different subjects (Figure 3A), representing the value of D as a function of the different frequencies over which the comparisons were performed, i.e.: in the range of 1 to 40 Hz for the EEG and 1 to 10 Hz for the PPG. As explained in the methods, SE1 was calculated within a window of size 5 Hz for each frequency value on the x-axis. Thus, the entropy was calculated in the range from 1 to 6 Hz for the value of 1 Hz on the x-axis, and in the range from 2 to 7 Hz for the value of 2, etc. Because the EEG and the PPG had different upper limits along the x-axis^2^, the Bonferroni significance threshold also differed for the two modalities, as 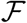 was equal to 40 for the EEG and 10 for the PPG.

**Fig. 2.**
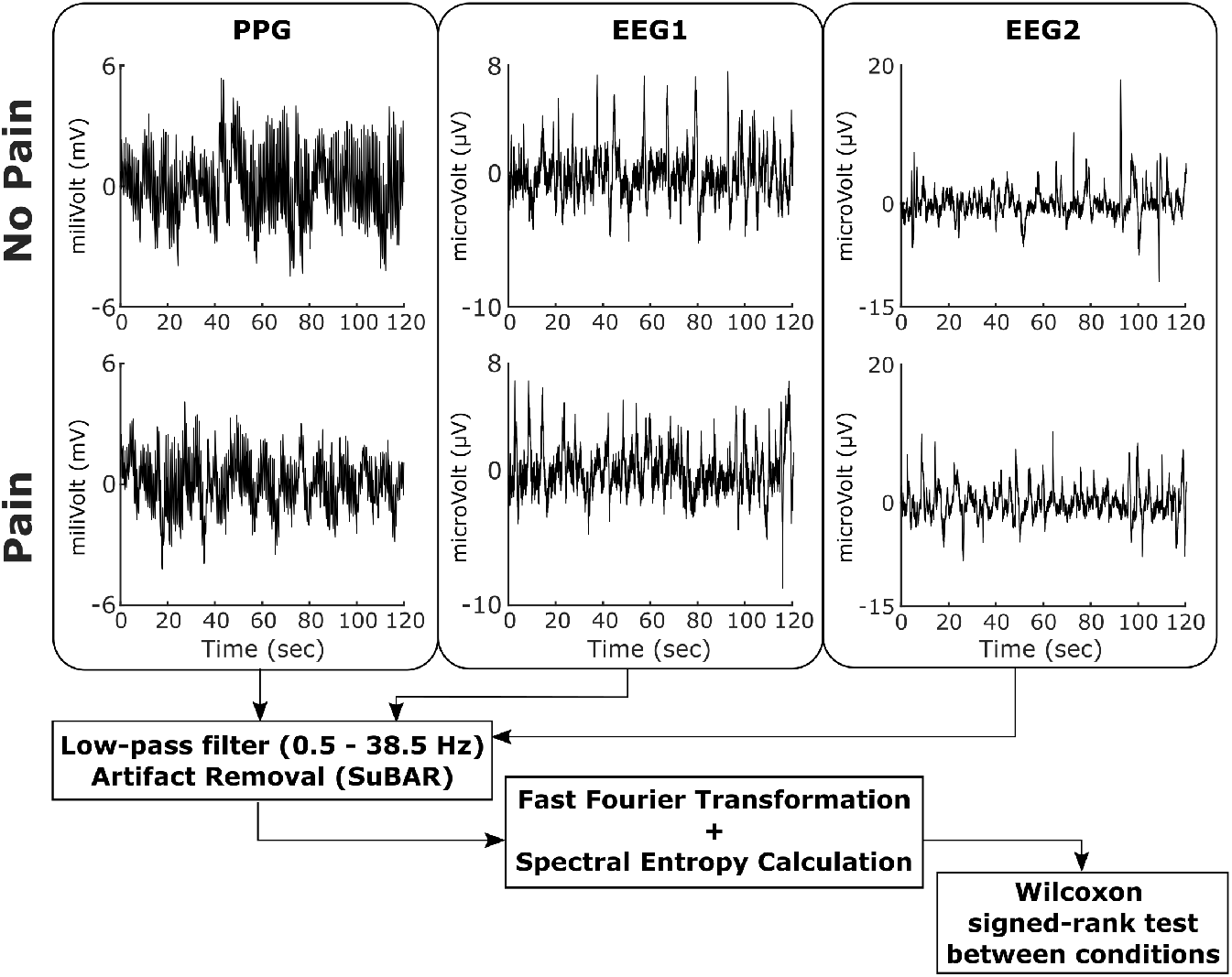
Signal processing workflow and statistical analysis. For the analysis, we first separated the signals belonging to the different conditions, pain or no pain, applying a low-pass filter and artifact removal to each signal. The *fft* MATLAB function was then used to calculate the power spectrum, which was used to assess the spectral entropy (SE) at different frequency bands and with frequency windows of 5 Hz. Finally, we compared the SE across different conditions over different frequency ranges and across different participants.

**Fig. 3.**
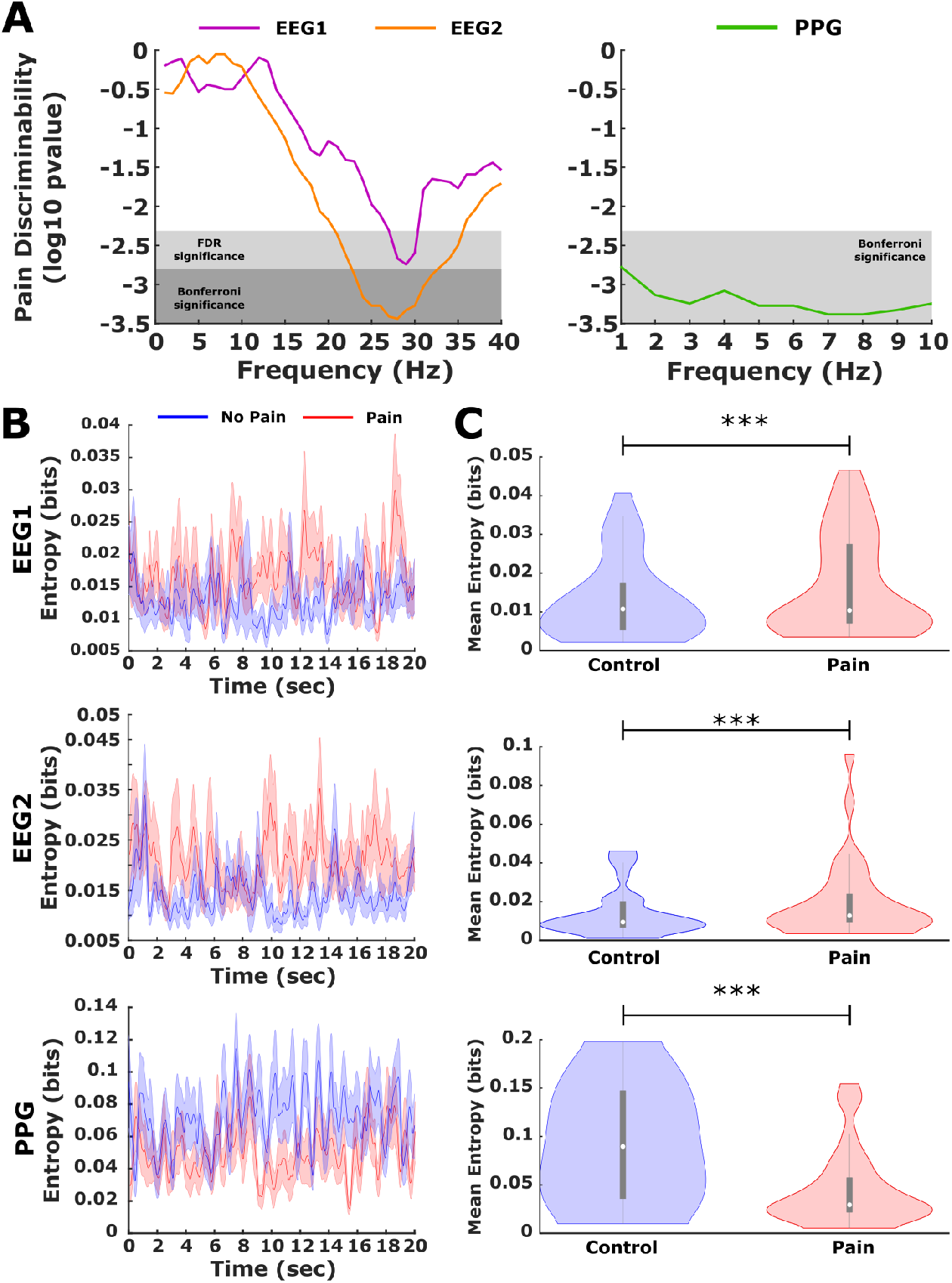
Discriminability of painful stimulation by 1D spectral entropy. **A**: Discriminability 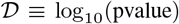 as a function of the different frequencies over which the conditions were compared for values of SE1 using a Wilcoxon signed-rank test (Methods). For each signal and condition, SE1 was calculated over the entire time series. The temporal average was then taken and the resulting mean values of SE1 compared between the two conditions. Frequencies range from 1 to 40 Hz for the EEG and from 1 to 10 Hz for the PPG. For each frequency value on the x-axis, SE1 was calculated within a window of 5 Hz. Thus, for the value of 1 Hz in the x-axis, entropy was calculated in the range 1-6 Hz and similarly, for a value of 2 Hz it is in the range from 2-7 Hz. The FDR region and Bonferroni correction significance is marked by light and dark gray rectangles, respectively. **B**: SE1 as a function of time for fixed frequency values (29 Hz for EEG1, 28 Hz for EEG2 and 7 Hz for PPG) over a time interval of 20 sec. **C**: After averaging the temporal signal of SE1 (illustrated in panel B), comparing the conditions highlighted significant differences between pain and no pain for all signals. The gray rectangles within the violins represent the first and third quartiles, and the white dot within those rectangles represents the median of the distributions: *** indicates p < 0.005.

SE1 was able to discriminate the pain condition for the three sets of sensory data, EEG1 (purple line), EEG2 (yellow line) and PPG (green line), with the major discriminability found at 25-30 Hz for the EEG and at 5-10 Hz for the PPG (the region of Bonferroni significance is illustrated in figure 3A with a transparent dark-gray rectangle, while that for FDR correction is marked in light-gray). The evolution of SE1 over time was obtained from the three signals during 20 sec, which corresponded to 10,000 time points for a sampling frequency of 500 HZ (figure 3B). After taking the temporal average of SE1, the mean entropy across participants provided significant differences between the conditions, as illustrated in figure 3C. The values of maximum discriminability of SE1 for the three classes of signals EEG1, EEG2 and PPG were achieved at 29 Hz (p< 10^-2^), 28 Hz (p< 10^-3^) and 7 Hz (p< 10^-3^), respectively.

We also assessed the discriminability between the conditions that achieved SE2, analyzing the 2D SE values at different frequencies for the pairs of signals (EEG1, EEG2), (EEG1, PPG) and (EEG2, PPG), as well as across different frequency ranges (figure 4). Like SE1, SE2 was also calculated in squared windows of size 5 × 5 Hz^2^. None of the comparisons of SE2 survived the Bonferroni correction or the FDR and thus, we only report here the uncorrected p-values. The values of maximum discriminability and the associated p-values occurred at (2,5) Hz for (PPG, EEG1: p< 10^-2^) and at (33,31) Hz for (EEG1, EEG2: p ≈ 0.01), yet they were not significant for PPG,EEG2 (p=0.05).

**Fig. 4.**
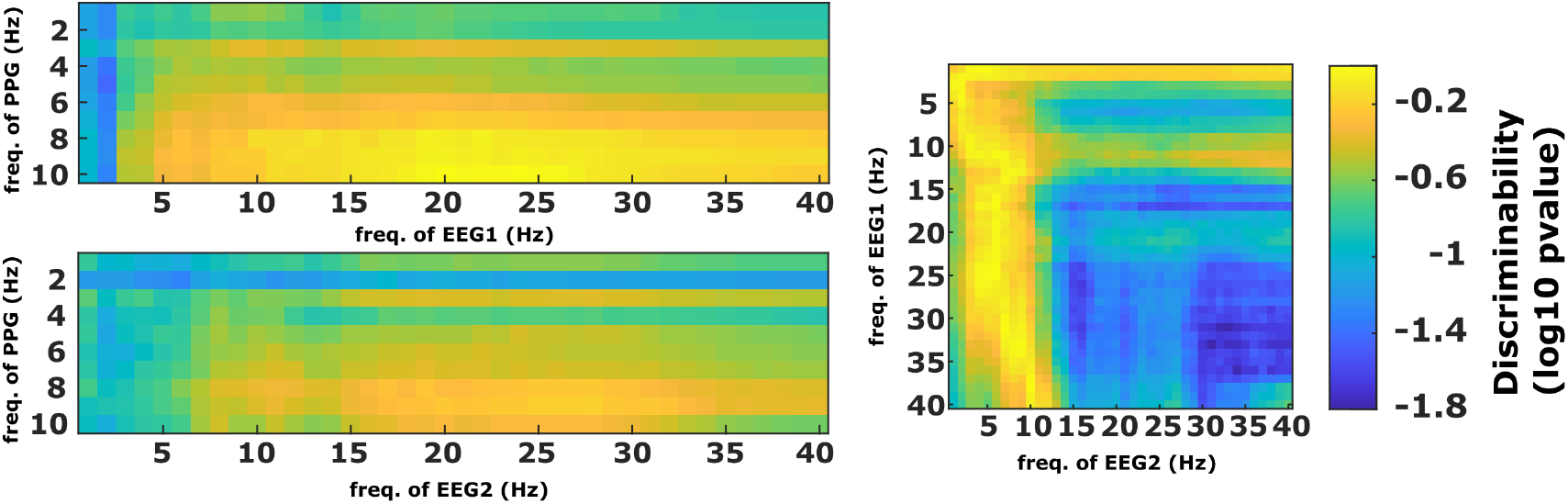
Discriminability of painful stimulation by 2D spectral entropy. A similar strategy was followed to the SE1 strategy (figure 3) but now calculating the 2D spectral entropy (SE2). After comparing the values of the temporal mean of SE2 matrix entries using a Wilcoxon signed-rank test for different conditions (pain vs no pain). The surfaces of discriminability achieved by pairs of different signals (EEG1, EEG2), (EEG1, PPG) and (EEG2, PPG) were represented across different frequency ranges. Similar to SE1, SE2 was also calculated in squared windows of 5×5 Hz^2^.

## IV. Discussion

Pain perception is a prototypic example of a well-orchestrated systemic response. Here, we have developed a compact device to simultaneously record physiological brain and heart parameters in response to a well-controlled painful thermal stimulus. The device consists of a Peltier cell that allows the temperature to be precisely varied under the control of an external computer, two EEG frontal electrodes (attached to the left and right hemispheres) and a PPG sensor located on one finger of the opposite hand to that on which the Peltier cell is placed. This platform can provide very precise information on the thresholds of maximum thermal tolerance to heat, which can potentially serve to assess novel strategies for both the diagnosis and follow-up of different pathological conditions.

Several studies have assessed the brain’s response to heat induced pain using EEG [22]–[24], yet the novelty of this study is that a compact setup has been developed to record the physiological responses of the heart and the brain while a subject receives a painful heat stimulus. This device enabled the mean heat pain threshold to be assessed in the cohort, defining a threshold of 42.68°C in a population with a mean age of 30-years-old. This result was consistent with previous studies using quantitative sensory testing (QST) [18], [21], thereby validating the reliability of our device to measure heat pain thresholds.

From a methodological point of view, it is important to note that these values were obtained when heating one of the subject’s hands for the first time. When we repeated the same procedure a second time on the contralateral hand soon after heating the other hand, the physiological responses in the PPG differed from that reported here (data not shown). Hence, heating one hand affected the physiological response of the subsequent heating of the contralateral hand. Accordingly, further studies will be needed to fully clarify these relationships as here we focused on signals from the initial heating of a *naive* hand, the right hand in 23 subjects and the left hand in the remaining 12 participants (for exact values see Table I).

Our study shows that thermal pain is characterized by a reduction in the entropy of the heart response measured by PPG, which suggests that in addition to the autonomic nervous system’s response, exposure to a thermal pain stimulus decreases the unpredictability of physiological systems as measured by their SE. In agreement with previous studies, we propose that upper supraspinal centers might fulfil a critical role in this physiological process [25]. Moreover, the posterior ventral nucleus of the thalamus might also coordinate the visceral sensitivity as it is activated by mechanical and thermal stimuli in a nociceptive range [26]. Furthermore, we suggest that the primary and the secondary somatosensorial cortex, in addition to the Anterior Cingulate Cortex might be involved in this physiological mechanism [27], [28].

Other algorithms based on SE in different frequency bands have been introduced to monitor physiological states. The most widely known is the bispectral index (BIS), which is used in daily clinical practice to monitor the depth of anesthesia during surgical interventions in real-time [29], [30]. Through a device that uses four EEG electrodes located on the patient’s forehead to measure the brain’s electrical activity, BIS calculates the SEs in different frequency bands and combines them using a proprietary algorithm to produce a numeric index between 100 (no anesthesia) and 0 (maximum anesthesia, where the level of consciousness measured by the frontal EEG activity is zero). The FDA have validated that BIS levels between 40 and 60 are adequate for general anesthesia during surgical interventions. For pain perception, such FDA-approved indices do not exist to date.

It is well-known that several chronic pain syndromes are associated with alterations to the activity of the Descending Nociceptive Inhibitory System (DNIS), such as fibromyalgia, painful diabetic neuropathy and lower back pain [31]–[33]. The DNIS is comprised of a network of cortical and subcortical brain areas, including the anterior insula, middle frontal gyri and amygdala, and the rostral ventromedial medulla and periaqueductal gray brainstem regions, which can inhibit nociceptive afferent brain input [34], [35]. A relationship between the DNIS and Heart Rate Variability (HRV) has been shown, whereby patients with an impaired DNIS have a lower resting HRV [36], consistent with the reduced entropy found here [37], [38]. Moreover, an abnormally low HRV was detected in a group of individuals with Chronic Fatigue Syndrome, indicating that they might have weaker parasympathetic modulation of their heart rate [39]. In other chronic pain syndromes like fibromyalgia, a reduced HRV was thought to reflect weaker emotional adaptability and resistance to stress [40].

Our findings reveal a close relationship between pain perception and the brain’s physiological entropy at different frequencies, in agreement with previous studies showing that pain is associated with a spatially extended network of dynamically recruited brain areas, resulting in complex temporal–spectral patterns of brain activity [41]. In particular, pain produces individual variations in the SE of both EEG and PPG at different frequencies (measured by SE1), as well as in their bi-dimensional interaction (as by SE2). Pain-related neuronal oscillations were observed previously at Theta (4–7 Hz), Alpha (8–13 Hz), Beta (14–29 Hz), and Gamma (30–200 Hz) frequencies [42]–[45]. Here, we found that both left-hemisphere EEG and right-hemisphere EEG have the best discrimination of the painful stimulus in the Beta and Theta band, and then in the Alpha and Gamma bands, as seen elsewhere [41], [42], [46]. EEG responses in the two hemispheres at frequencies around 28 Hz were seen to discriminate the painful state, in agreement with a previous assessment of the adequacy of analgesia in which such oscillations were shown to participate in discriminating painful events [47].

Our study has also some limitations. First, although we tried to keep the volunteers calm by playing relaxing music while recording the physiological response to the thermoalgesic stimulus, we did not control the arousal, attention or salience, nor did we assess cognitive appraisal before or during the experiment. Second, we focused this study on thermoalgesic stimulation, yet different painful stimuli could be incorporated into our device for future studies, for instance mechanical pain, providing greater sensitivity and specificity to discriminate different classes of painful stimulation. Finally, the temperature of the Peltier cell was varied in increments of 0.5°C to achieve the activation of C receptors (responsible for heat processing) together with that of the A*δ* fiber receptors (responsible for noxious processing stimuli) [19], [20], a critical constraint to our design. However, this protocol did not allow the thermoalgesic exposure to heat to be randomized, which could possibly be incorporated into the stimulation protocol in future studies.

In summary, our compact device allows brain and heart physiological signals to be recorded simultaneously in response to well-controlled thermal pain stimuli. We show that the SE of the physiological signals can discriminate pain states. Future work should validate similar metrics based on SE for the on-line variation of painful stimuli, or the dynamic on-line interaction between PPG and EEG signals, for instance using Granger causality [48]–[51] or transfer entropy [52], [53], as used previously to establish different dynamic brain mechanisms in pain-related conditions like migraine [54], [55]. Last but not least, future studies should assess whether our dual EEG and PPG system is useful to study some pathological conditions in which the autonomic nervous system functions abnormally, such as small fiber neuropathies, fibromyalgia or painful diabetic neuropathy.

## Acknowledgements

We acknowledge the assistance of Javier Rasero through critical reading of this manuscript and his constructive suggestions for the statistical analysis. JMC is funded by Ikerbasque: The Basque Foundation for Science, the Ministerio Economia, Industria y Competitividad, Spain and FEDER (grant no. DPI2016-79874-R), and the Department of Economic Development and Infrastructure of the Basque Country (Elkartek Program, KK-2018/00032 and KK-2018/00090). AE is funded by the ELSC and by the Department of Education of the Basque Country postdoctoral program (POS-2019-2-0020). EMG is funded by the Department of Education of the Basque Country postdoctoral program (POS-2019-1-0034).

## Author contributions

Manufacturing of the hardware device: MNI, JOAC; Patient recruitment: MNI; Design and implementation of the signal processing methods: BCP, ID, AE, JMC; Figure preparation: MNI, BCP, JMC; Draft of the manuscript: MNI, BCP, ID, AE, EMG, SS, JOAC, JMC; Supervision of the research - equal contribution: JOAC, JMC;

---

**Fig. S1.**
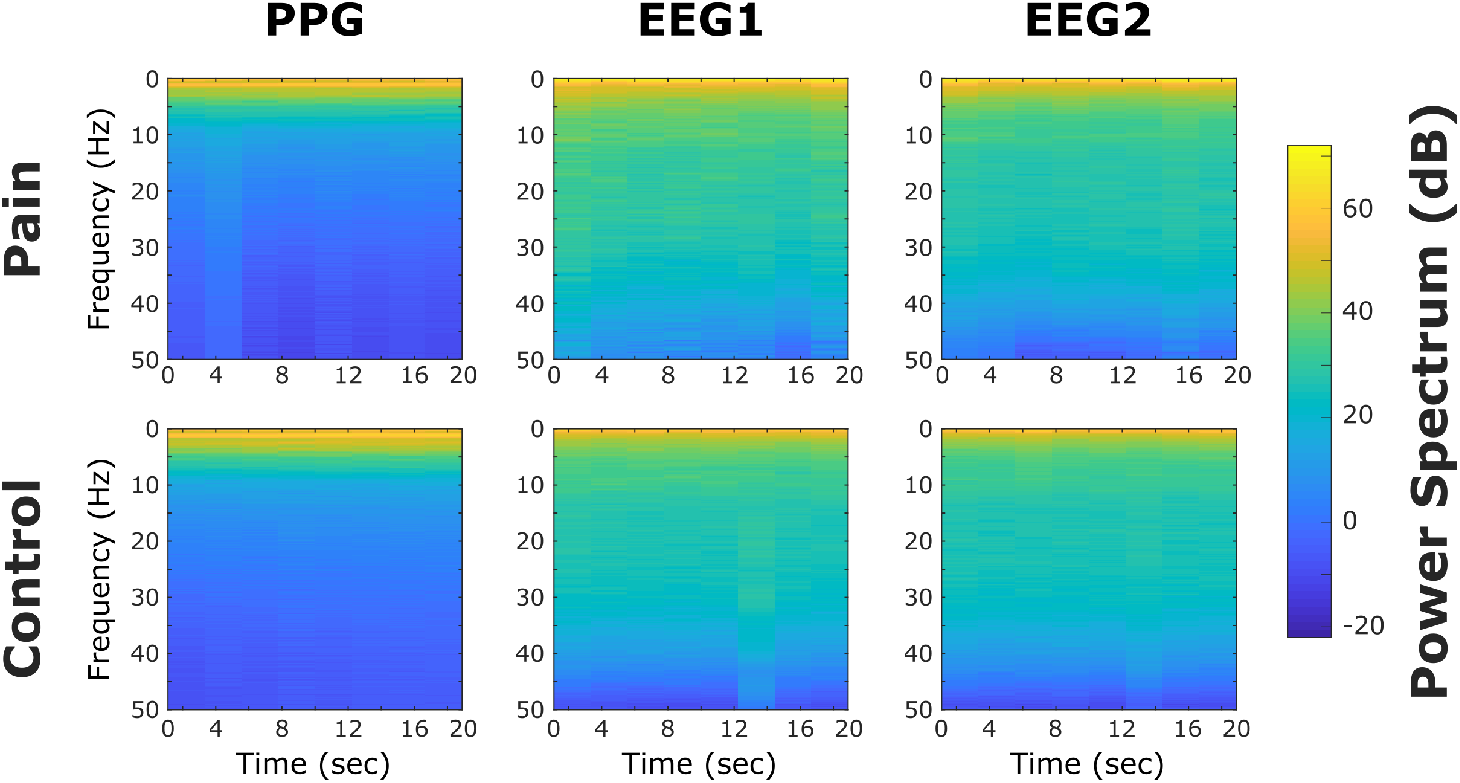
Population spectrograms of the different signals across conditions. Population spectrograms were obtained by simply averaging individual spectrograms from all the participants. The power spectrums are represented in units of decibels (dB) by simply calculating the transformation of 20 log_10_(S), where S is the mean value of all the subjects of the complex modulus of a short-time Fourier transform of the signal. From these spectrograms, we chose an upper limit of 40 Hz for the EEG and 10 Hz for the PPG. These limits defined the maximum value of frequencies used to calculate SE1 and SE2, and therefore, to compare SE1 and SE2 between the conditions of pain and no-pain.

## Footnotes

1 PPG signals provide an indirect measure of the heart’s physiological response. When the variability in the heart rate associated with the signal is assessed, both PPG and a direct measure like an electrocardiogram (ECG) provide very similar values, which makes PPG a good proxy to study dynamic the variations in the response of the heart.

2 The upper limits of 40 Hz for the EEG and 10 Hz for the PPG were estimated by examining their corresponding spectrograms for the conditions (figure S1), simply choosing a value at which the spectrum was negligible for frequency values above the chosen limit.

